# Catch & Release – rapid cost-effective protein purification from plants using a DIY GFP-Trap-protease approach

**DOI:** 10.1101/2025.05.15.654187

**Authors:** Sebastian Schwartz, Carina Engstler, Susanne Mühlbauer, Eduard Windenbach, Tobias Wunder, Hans-Henning Kunz, Benjamin Brandt

**Author notes:** corresponding author: Benjamin Brandt. senior authors: Hans-Henning Kunz and Benjamin Brandt.

## Abstract

The purification of proteins is the foundation to study their structure, function, biochemical properties, and interaction partners. In plant research, unique challenges arise from the complexity of plant tissues, interference of secondary metabolites, and sometimes the low abundance of target proteins. Many conventional plant protein purification methods rely on expensive reagents, multi-step procedures, and labor-intensive workflows, limiting their feasibility for many applications. Here, we present the “Catch & Release” system, a cost-effective, fast and reliable one-step purification workflow for the isolation of soluble and membrane-bound proteins from plant tissues. The Catch & Release toolbox includes a vector set, a homemade GFP-trap and homemade proteases. Catch & Release vectors streamline cloning and transgenic plant selection through the Fluorescence-Accumulating Seed Technology (FAST), which marks positive transformants with a strongly fluorescing seed coat. Each plasmid consists of four, easy to exchange, modules: a plant promoter, a cloning dropout marker, protease cleavage sites, and seven different epitope tags, including an innovative dual-fluorescent tag, providing flexibility for diverse experimental needs. The *in vivo* functionality of all modules has been confirmed. Besides enabling standard molecular biological experimentation, our vector set in combination with homemade GFP-trap and proteases enables efficient and rapid isolation of soluble and high molecular weight membrane proteins directly from plants. By following our detailed reagent preparation instructions, purification costs can be decreased hundred-fold compared to the commercially available options.

## Introduction

Understanding the function, structure, and regulation of proteins is fundamental to deciphering complex biological processes. However, as the biochemistry textbook states: “Never waste pure thoughts on an impure protein” (Berg et al., 2019). Consequently, the ability to isolate and purify specific proteins in an active and native state is a cornerstone of plant molecular biology research (Amrhein et al., 2015). Purifying proteins from plant sources presents significant challenges compared to microbial or mammalian systems. The inherent complexity of plant tissues, often rich in cell walls, vacuoles, and interfering secondary metabolites (phenolics, pigments, polysaccharides) complicates extraction (Schillberg et al., 2019) and can compromise protein stability and activity (Du et al., 2022). Furthermore, many plant proteins of interest are expressed at low endogenous levels requiring sensitive and efficient isolation strategies (Righetti and Boschetti, 2020).

The requirement for highly pure protein isolates is especially apparent in structural biology. Due to the above-mentioned limitations, most plant protein expressions and isolations are carried out in heterologous systems, primarily *E. coli.* In fact, 79% of high-quality protein structures from Arabidopsis proteins found in the protein data bank (PDB; https://www.rcsb.org) were obtained using *E.coli*-expressed proteins. However, heterologous expression of plant proteins, especially in prokaryotes, has considerable limitations such as the absence of eukaryotic post-translational modifications (PTMs) (Zhang et al., 2022), the lack of specific plant chaperones or cofactors and a non-native intracellular redox state (Bhatwa et al., 2021), all of which may affect correct folding. Improperly folded proteins can potentially yield structural or functional data not accurately reflecting the native state. These missing factors in combination with disparities in codon usage affecting translation fidelity and speed (Gustafsson et al., 2004) also frequently result in plant proteins not expressing in prokaryotes. To overcome these limitations, the expression of plant proteins in their native expression host is favorable.

Conventional protein purification workflows often rely on time consuming, multi-step chromatographic procedures, using expensive commercial affinity resins and reagents. Affinity tagging strategies (e.g., His-tag or Strep-tag) are commonly used for rapid-one-step isolation of recombinant proteins from many expression systems. However, their usability in plants is limited. This is in part due to the generation of stable transgenic lines expressing tagged proteins, which, depending on the species, can be a time-consuming process. Another reason is secondary metabolites interference with the tag-resin binding. These limitations are demonstrated by only 2% of Arabidopsis protein structures deposited in the PDB originate from protein isolated from Arabidopsis, all of which using complex multi-step purification methods. These combined factors – tissue complexity, low abundance and no suitable one-step affinity tags resulting in lengthy workflows – limit the throughput, scalability, and accessibility of protein purification from plants.

Advanced modular cloning systems, such as those based on Golden Gate assembly utilizing Type IIS restriction enzymes (e.g., BsaI, BpiI), have been developed for the construction of complex expression vectors (Engler et al., 2008). These systems enable the ordered, assembly of multiple DNA fragments (promoters, coding sequences, tags, terminators) in a single reaction which can be subsequently assembled into multi-expression unit constructs in a second reaction. However, establishing and utilizing Golden Gate or related plant-specific systems like GreenGate (Lampropoulos et al., 2013) requires a significant upfront investment including creating or acquiring libraries of standardized and compatible DNA modules. During the creation of sequence-verified entry modules the DNA sequences must be free of the specific Type IIS recognition sites used for the subsequent assemblies. This can be challenging, especially for large or complex DNA fragments such as genomic plant DNA fragments. While powerful for multi-expression unit constructs, the complexity and setup effort associated with these systems might be less justified for the routine generation of simpler vectors commonly used for expression and purification of tagged proteins.

To resolve current limitations for the purification of plant proteins in their native hosts, we developed and combined publicly available resources into the “Catch & Release” system, a versatile and cost-effective toolbox designed to enable efficient protein expression and purification of plant proteins. This system integrates several key features: (i) efficient vector construction using multiple cloning strategies facilitated by a visual dropout marker; (ii) accelerated generation of stable transgenic lines through integration of Fluorescence-Accumulating Seed Technology (FAST) (Shimada et al., 2010); (iii) a set of seven interchangeable C-terminal tags, including standard small peptide tags, fluorescent proteins, and innovative dual-fluorescent tags, providing flexibility for detection, localization, and purification; (iv) fluorescent tags in combination with protease cleavage sites (TEV or HRV 3C), which are compatible with homemade proteases and a DIY GFP-Trap to enable cheap and rapid one-step purification of native target proteins expressed in plants.

In this work, we demonstrate the utility and robustness of the Catch & Release toolbox for both transiently (*Nicotiana benthamiana*) and stably (*Arabidopsis thaliana*) expressed plant proteins. We show its successful application for the functional complementation, correct subcellular localization, and efficient purification of POTASSIUM CATION EFFLUX ANTIPORTER 1 (KEA1), a ∼ 123 kDa large, chloroplast inner envelope membrane protein. Furthermore, we validate the system for purifying the soluble stromal enzyme PHOSPHOGLYCERATE DEHYDROGENASE 3 (PGDH3) as intact (dimeric) and enzymatically active protein using the novel dual-fluorescent tag.

## Results and discussion

### Catch & Release includes a flexible vector set for cloning and protein analysis

To overcome common bottlenecks in plant transformation and functional protein studies, we established the “Catch & Release” toolbox (Fig. 1). The binary vector system included in the toolbox leverages a modular vector design, accelerating construct generation, simplifying transgenic line selection, and providing flexibility for downstream protein characterization.

**Figure 1.**
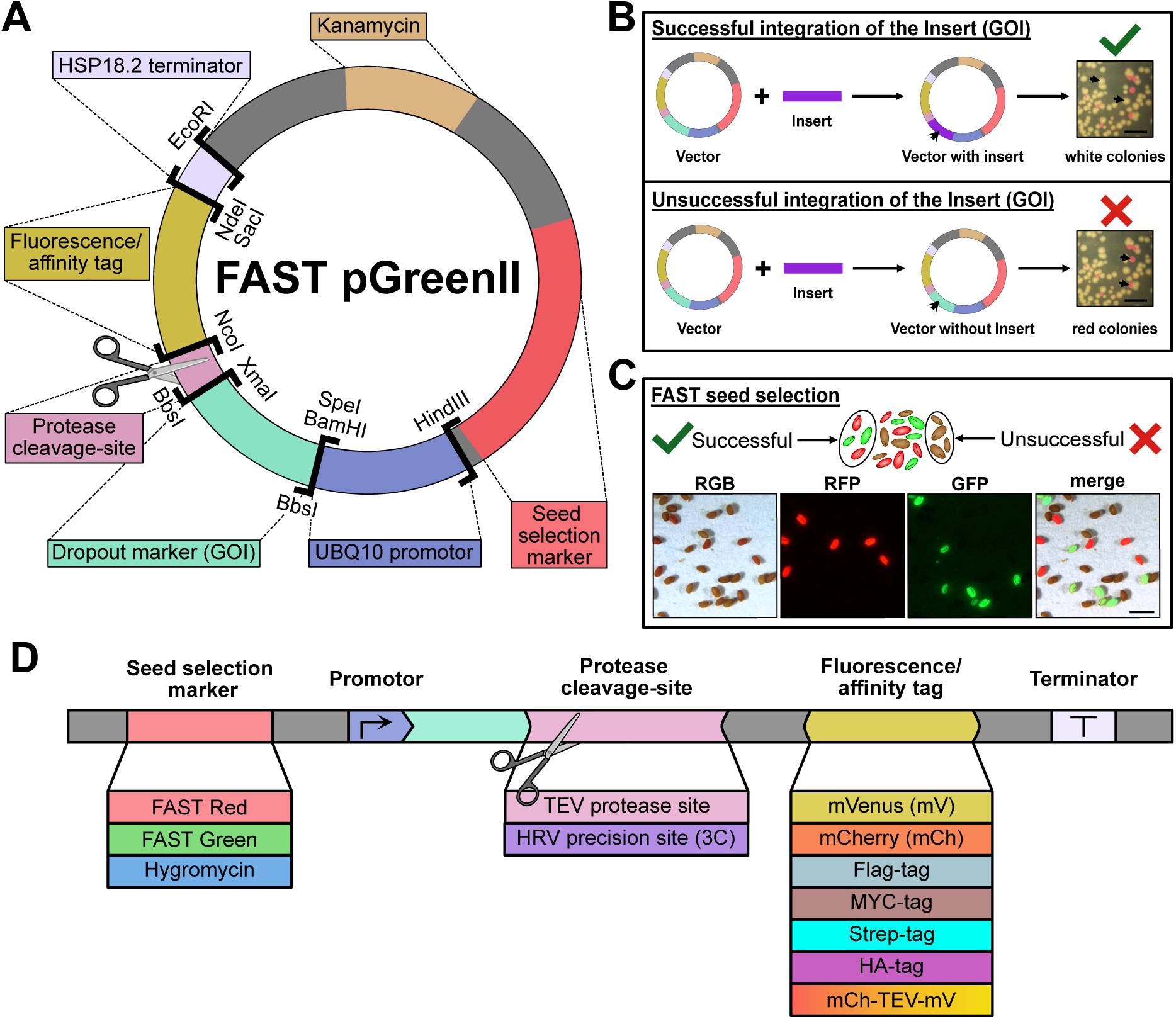
Overview and features of the Catch and Release vector set. (**A**) General design of the Catch and Release binary vectors highlighting key restriction sites (**B**) Red/white colony screening for successful DNA incorporation via Golden Gate-, Gibson- or classical restriction cloning. Scale bar = 0.5 cm (**C**) Seeds from Arabidopsis plants transformed with FAST-Green and FAST-Red vectors, examined under a fluorescence stereoscope. Strongly fluorescent seeds were selected for propagation to obtain homozygous T_3_ lines. Scale bar = 1 mm (**D**) Catch and Release starter set comprising 13 parts, categorized into four subtypes with a total of 30 vectors (listed in tab. 1). Vectors incorporating the HRV 3C precision site are available exclusively with the mVenus tag and the dual-fluorescent tag.

**Table 1:** Catch & release vector set including Addgene IDs.

| Construct | Addgene ID |
| --- | --- |
| pG20_Hyg | t.b.a. |
| pG20_TEV_mVenus_Hyg | t.b.a. |
| pG20_TEV_mCherry_Hyg | t.b.a. |
| pG20_mCherry_TEV_mVenus_Hyg | t.b.a. |
| pG20_3C_mCherry_TEV_mVenus_Hyg | t.b.a. |
| pG20_TEV_2xStrepII_Hyg | t.b.a. |
| pG20_TEV_MYC_Hyg | t.b.a. |
| pG20_TEV_HA_Hyg | t.b.a. |
| pG20_TEV_Flag_Hyg | t.b.a. |
| pG20_3C_mVenus_Hyg | t.b.a. |
| pG20_FR | t.b.a. |
| pG20_TEV_mVenus_FR | t.b.a. |
| pG20_TEV_mCherry_FR | t.b.a. |
| pG20_mCherry_TEV_mVenus_FR | t.b.a. |
| pG20_3C_mCherry_TEV_mVenus_FR | t.b.a. |
| pG20_TEV_2xStrepII_FR | t.b.a. |
| pG20_TEV_MYC_FR | t.b.a. |
| pG20_TEV_HA_FR | t.b.a. |
| pG20_TEV_Flag_FR | t.b.a. |
| pG20_3C_mVenus_FR | t.b.a. |
| pG20_FG | t.b.a. |
| pG20_TEV_mVenus_FG | t.b.a. |
| pG20_TEV_mCherry_FG | t.b.a. |
| pG20_mCherry_TEV_mVenus_FG | t.b.a. |
| pG20_3C_mCherry_TEV_mVenus_FG | t.b.a. |
| pG20_TEV_2xStrepII_FG | t.b.a. |
| pG20_TEV_MYC_FG | t.b.a. |
| pG20_TEV_HA_FG | t.b.a. |
| pG20_TEV_Flag_FG | t.b.a. |
| pG20_3C_mVenus_FG | t.b.a. |

“Catch & Release” plasmids offer cloning versatility, since they are fully compatible with most common methods such as GoldenGate assembly (Engler et al., 2008), Gibson/InFusion (Gibson et al., 2009) assembly, SLIC (Jeong et al., 2012) and traditional restriction-ligation cloning. Facilitating efficient screening during the cloning procedure, the vectors possess a visual dropout marker cassette containing a constitutively expressing red fluorescent protein (RFP) gene (Fig. 1A). This feature enables the identification of recombinant clones directly on the Kanamycin supplemented selection plate (Fig. 1B). Colonies with successfully incorporated DNA insert loose RFP expression and appear white. In contrast, non-recombinant background colonies remain red. This visual screening streamlines cloning by reducing laborious PCR or digestion checks.

To accelerate the generation of homozygous transgenic plant lines, a major time constraint in plant research, we incorporated the Fluorescence-Accumulating Seed Technology (FAST) (Shimada et al., 2010) into the vector set. The FAST system enables non-invasive selection of transformed seeds based on seed coat fluorescence, eliminating the need for antibiotic or herbicide resistance markers that can impair plant development (Yin et al., 2008) and trigger pleiotropic phenotypic alterations. Additionally, using fluorescent seed coats as transgenic marker result in major time savings during segregation analyses and establishment of homozygous plant lines. Our toolbox includes both FAST-Red and FAST-Green options for fluorescence-based seed sorting thus enabling selection for plants carrying multiple transgenes (Fig. 1C). In addition, conventional Hygromycin-based selection vectors are also available to accommodate for a variety of experimental requirements. All section markers included in the vector set work side-by-side with the most common selection markers used in the major seed collections (BASTA, Kanamycin, sulfadiazine).

A cornerstone advantage of the Catch & Release toolbox lies in its variable protein tagging strategy (Fig. 1D), which offers a suite of seven distinct C-terminal tags, encompassing small peptide tags (e.g., Strep, Flag, MYC, HA) for detection and isolation and single fluorescent proteins (e.g., mVenus, mCherry) for localization, isolation, and detection. In addition, we designed a novel dual-fluorescent tag, which is offered in two configurations that differ in mCherry’s removability: mCherry-TEV-mVenus (non-removable mCherry) and 3C-mCherry-TEV-mVenus (fully removable mCherry). Critically, each tag is preceded by protease cleavage sites (TEV or HRV 3C) engineered to ensure tag removal leaving only a minimal number of amino acid residues behind on the target protein. Removable tags permit initial purification/detection followed by controlled cleavage *in vitro* to release the native protein. This approach ensures the suitability of the protein for further downstream functional and structural analyses by preventing potential interference with the fusion tag.

The system’s modularity extends beyond tags. The set comprises 13 exchangeable components, yielding 30 ready-to-use binary vectors (Tab. 1). Importantly, key functional elements like protease sites, promoters and terminators (e.g., the commonly used UBQ10 promoter and HSP18.2 terminator) (Pratt et al., 2020) can be readily exchanged using designated restriction sites (Fig. 1A). This adaptability empowers researchers to easily customize vectors for specific expression requirements such as tissue-specific expression, or usability in different plant species.

### Functional validation of Catch & Release constructs in transgenic plants

To validate the functionality of proteins expressed using our vectors, we used our most bulky tags to assess their risk of interference with protein function. Here, we performed complementation studies in the *Arabidopsis thaliana kea1-1kea2-1* double mutant (Kunz et al., 2014). This loss of function mutant lacks the plastid inner envelope cation efflux antiporters KEA1/KEA2, which results in delayed chloroplast development and impaired photosystem II quantum yield (*F*_v_*/F*_m_) in younger leaves (DeTar et al., 2021) (Fig. 2A-B).

**Figure 2.**
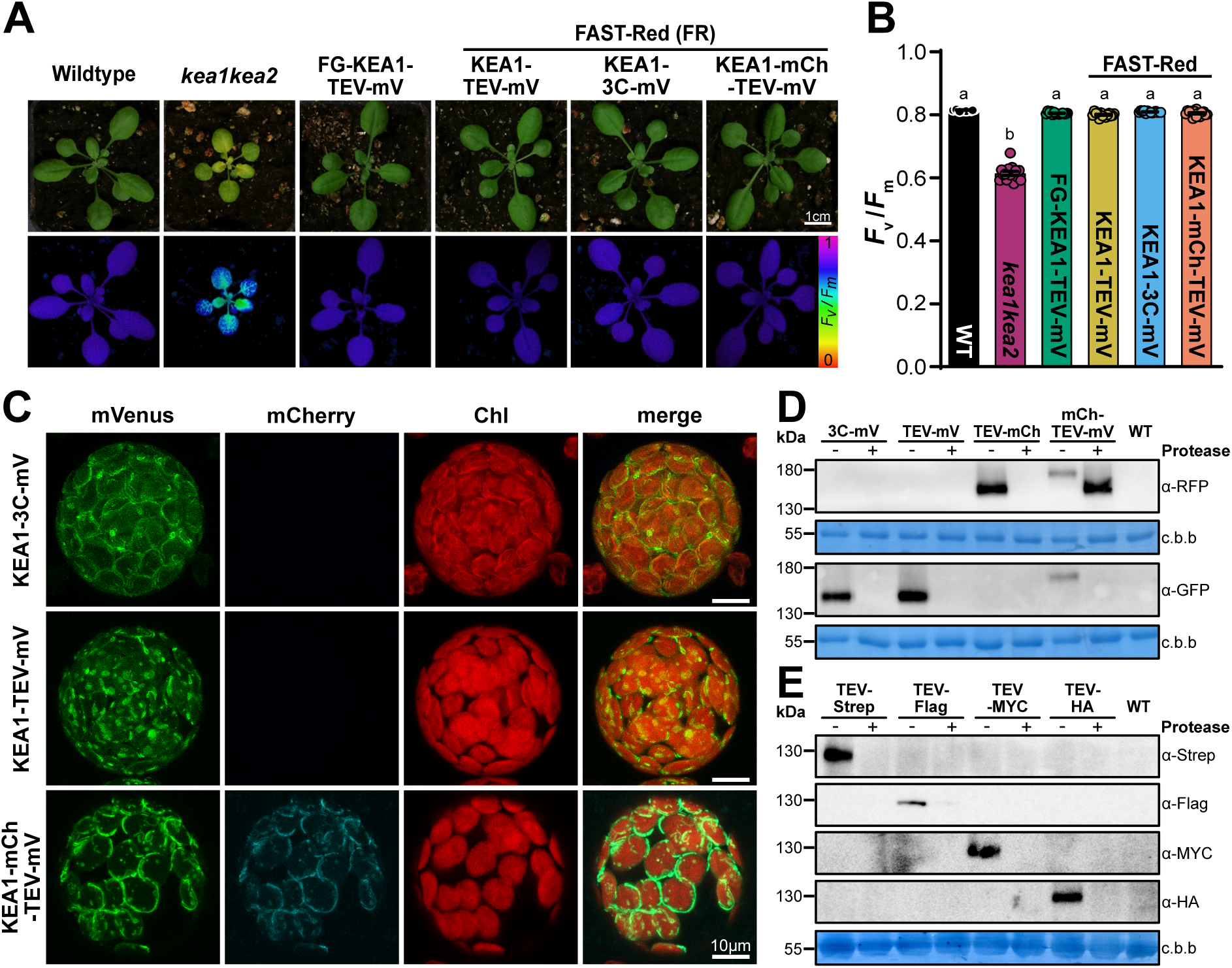
Functional characterization of KEA1 constructs using Catch and Release Vectors. (**A**) Photographs and PAM images (*F*_v_/*F*_m_) comparing WT, *kea1-1kea2-1*, and complemented *kea1-1kea2-1* lines expressing FAST-Green (FG)-KEA1-TEV-mVenus, FAST-Red (FR)-KEA1-TEV-mVenus, FAST-Red-KEA1-3C-mVenus, and FAST-Red-KEA1-mCherry-TEV-mVenus. Scale bar = 1 cm (**B**) *F*_v_/*F*_m_ in WT, *kea1-1kea2-1*, and complemented plants expressing FAST-Green(FG)-KEA1-TEV-mVenus, FAST-Red-KEA1-TEV-mVenus, FAST-Red-KEA1-3C-mVenus, and FAST-Red-KEA1-mCherry-TEV-mVenus. Data are presented as mean ± SEM (n = 14). Statistical significance was determined via one-way ANOVA, with different letters indicating significantly different groups. (**C**) Confocal laser micrographs of Arabidopsis protoplasts expressing KEA1-TEV-mVenus, KEA1-3C-mVenus, and KEA1-mCherry-TEV-mVenus. Fluorescence images show KEA1-mVenus (green, left), KEA1-mCherry (cyan, middle left), chlorophyll auto-fluorescence (red, middle right), and a merged view (right). (**D**) Immunoblot analysis of protease site accessibility in whole-leaf extracts from WT and complemented *kea1kea2* plants expressing KEA1-3C-mVenus, KEA1-TEV-mVenus, KEA1-TEV-mCherry, and KEA1-mCherry-TEV-mVenus. The same membrane was stained with coomassie brilliant blue (c.b.b.) as a loading control. (**E**) Immunoblot analysis of protease site accessibility in *N. benthamiana* leaves infiltrated with KEA1-TEV-Strep, KEA1-TEV-Flag, KEA1-TEV-MYC, and KEA1-TEV-HA. Un-infiltrated leaves served as WT controls. C.b.b. staining was used as a loading control.

We generated transgenic lines expressing KEA1 fused to C-terminal mVenus (via TEV or HRV 3C linker) or the dual mCherry-TEV-mVenus tag. Homozygous third generation progeny (T_3_) were selected efficiently using integrated FAST markers. Pulse-Modulated Chlorophyll Fluorescence measurements revealed that the reduced *F*_v_*/F*_m_ phenotype of the *kea1-1kea2-1* mutant (0.58 ± 0.03) was fully restored to wild-type levels (Col-0: 0.81 ± 0.01) in all complemented lines expressing tagged KEA1 variants (e.g., KEA1-TEV-mVenus: 0.81 ± 0.01) (Fig. 2A-B).

Next, we examined the localization of the tagged KEA1 proteins in protoplasts isolated from the complemented Arabidopsis lines. Confocal laser microscopy revealed that fluorescence signals from mVenus, or mCherry and mVenus (in dual-tagged lines) were localized to the periphery of chloroplasts (Fig. 2C), consistent with the known chloroplast envelope localization of KEA1 (Bölter et al., 2020). The precise co-localization of mCherry and mVenus signals in the dual-tagged construct further confirmed the integrity and proper trafficking of the fusion protein, demonstrating the system’s reliability for accurate subcellular localization studies, even with large fusion tags such as the mCherry-TEV-mVenus (54.7 kDa).

These results demonstrate that KEA1 proteins tagged using the Catch & Release toolbox are correctly expressed, folded, localized, and biologically active in planta, capable of rescuing physiological impairments. The successful application to KEA1 which presents considerable challenges due to its size, import into plastids and targeting to the inner envelope membrane, strongly suggests the system’s applicability to proteins with less complex folding or targeting. It also validates the system’s suitability for functional genetic studies.

We next aimed to confirm the accessibility of the protease cleavage sites, a central design principle. We tested protease assisted removal of the fluorescence tag fused to KEA1 by immunoblotting of total protein extracts from the stable transformed Arabidopsis lines (Fig. 2D). Antibodies against GFP (also detecting mVenus) and mCherry recognized proteins of the molecular weights consistent with the expected size of the KEA1 fusion proteins (∼150 kDa for single tags, ∼175 kDa for the dual tag). TEV and HRV 3C proteases were produced in house using publicly available expression vectors (Fig. S1A-D; see material and methods). Upon incubation of the extract with the appropriate protease (TEV or HRV 3C) for up to 4 hours, no immune signal for full length KEA1 fluorescent fusion proteins was detected indicative of protease-assisted removal of the tag. Treatment of the dual-tagged KEA1-mCherry-TEV-mVenus protein extract with TEV protease resulted in a clear shift in the mCherry signals apparent molecular weight and loss of the mVenus immunoblotting signal, confirming precise cleavage at the engineered protease site.

We also tested the expression, tag accessibility and removal using small peptide-tagged KEA1 constructs transiently expressed in *N. benthamiana* leaves (Fig. 2E), a widely used system for transient protein expression in plants. Immunoblotting detected proteins around 130 kDa, consistent with the expected size of KEA1 (∼123 kDa) (Bölter et al., 2020) plus the small peptide tags (e.g., Strep, Flag, MYC, HA). Subsequent treatment with TEV protease removed these tags as apparent by the loss of tag detection, confirming efficient tag cleavage in this transient *in planta* expression system.

These results not only validate the functionality of the small peptide tags and proteases assisted tag-removal but also showcase the Catch & Release toolbox’s adaptability across different plant species and expression strategies (stable vs. transient).

### Workflow for the isolation of native KEA1

The successful phenotypic complementation of the *kea1-1kea2-1* double mutant by the KEA1-TEV-mVenus construct served as the initial validation, confirming that C-terminal tagging with TEV-cleavable mVenus did not compromise the function or correct subcellular localization of the KEA1 transporter (Fig. 2A, 2C). This prompted us to use these lines for the purification of the large multi-membrane spanning plastid inner envelope protein KEA1. The presence of the TEV protease cleavage site engineered between KEA1 and the mVenus tag allows for specific affinity capture via a GFP-trap followed by its precise removal to yield native protein (Fig. 3A).

**Figure 3.**
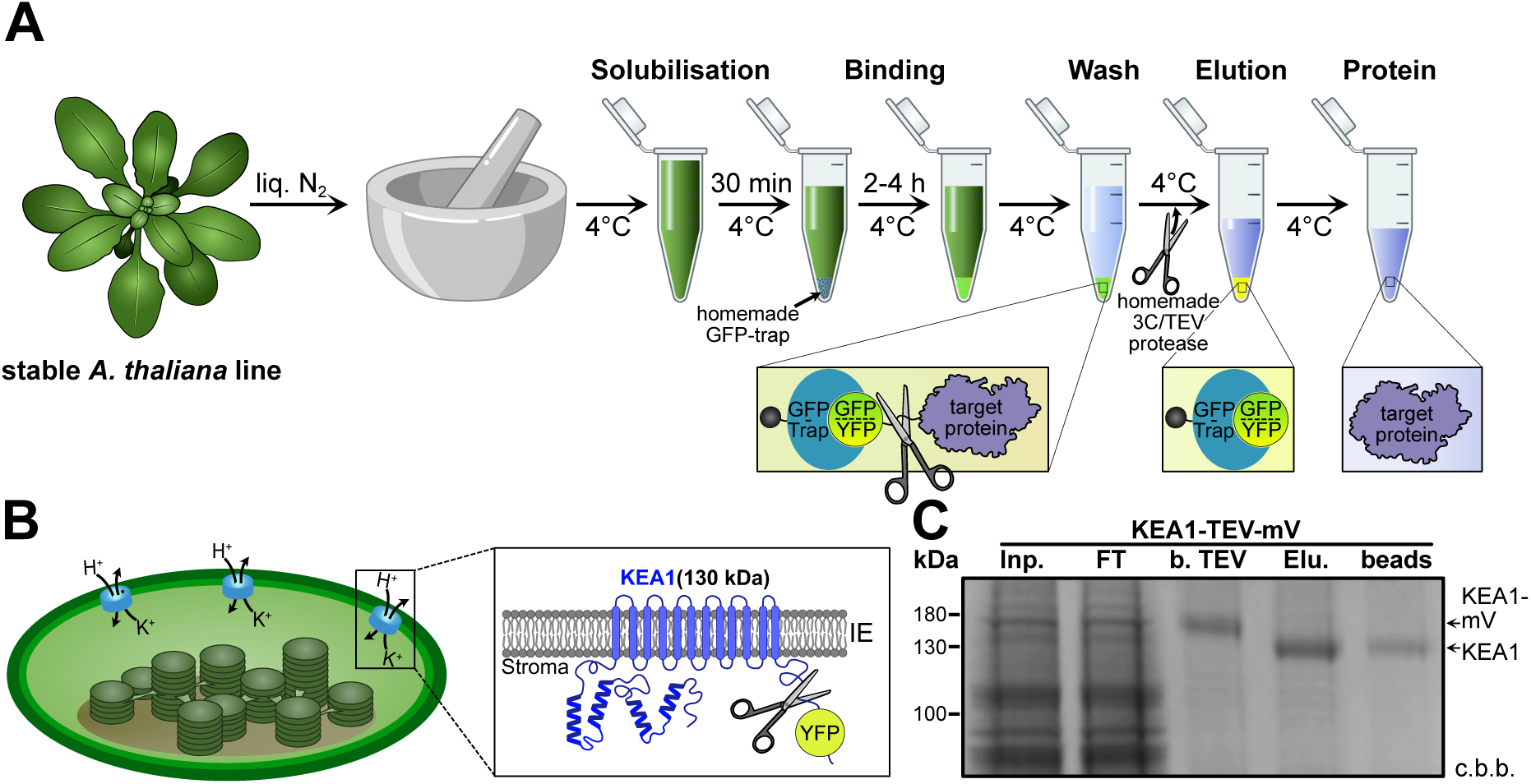
Isolation and structural features of KEA1 from *A. thaliana*. **(A**) Schematic overview of the protein purification workflow using the homemade GFP-trap and proteases. *A. thaliana* plants were first ground to powder in liquid nitrogen and solubilized for 30 minutes. The solubilized protein mixture was then incubated with homemade GFP-trap for 2-4 hours at 4°C, followed by multiple wash steps. Elution was triggered by protease digestion, resulting in the release of non-tagged protein. (**B**) Schematic representation of the plastid inner envelope potassium cation efflux antiporter KEA1. The N-terminal region is predicted to contain 2-3 coiled-coil domains (depicted as helical structures), while the C-terminal domain, facing the stroma, is fluorescently tagged and includes the KTN domain. (**C**) SDS-PAGE analysis verifying successful KEA1 purification from stable A. thaliana lines, using the described workflow. Lanes represent: Total protein lysate (Input, Inp.), unbound fraction after incubation with GFP-Trap (Flow-Through, FT), affinity-bound protein before protease cleavage (b. TEV), protein eluted after TEV cleavage (Elution, Elu.), and the GFP-Trap resin after elution (beads).

Building upon the validated KEA1-TEV-mVenus line, we developed and implemented a cost-effective and scalable one-step affinity purification strategy (Fig. 3A). Purifying KEA1, an integral membrane protein with up to 12 predicted transmembrane domains localized to the plastid inner envelope, presents significant biochemical challenges. Topological predictions place the N-terminus, containing putative coiled-coil domains, and the C-terminus, featuring the conserved KTN (K^+^ transport, nucleotide-binding (Roosild et al., 2002)) module, within the plastid stroma (Fig. 3B). The workflow centers on the use of a homemade GFP-trap affinity resin. The GFP binding DARPin of the homemade GFP-trap binds to GFP, YFP, CFP and their derivates (i.e. mVenus) with high affinity while no interaction could be detected to mCherry or mRuby (Hansen et al., 2017). Employing a slightly adapted protein purification and coupling strategy (Fig. S2 see material and methods for details), we were able to produce home-made GFP-trap currently worth approximately 30.000 Euro of commercially available products in only three working days. Combined with in-house produced TEV or HRV 3C proteases (Fig. S1A-B), this approach substantially lowers the financial barrier for protein purifications using a GFP-trap from plant tissues.

The protocol commenced with cryogenic tissue disruption to preserve protein integrity, followed by solubilization (Fig. 3A). Incubation with the GFP-trap resin under controlled conditions (4°C) facilitated efficient capture of the KEA1-TEV-mVenus fusion protein. SDS-PAGE analysis confirmed the isolation of the full-length KEA1-TEV-mVenus fusion protein, which migrated at an apparent molecular weight of approximately 150 kDa (Fig. 3C). Due to the high affinity between the GFP-Trap binding moiety and the mVenus tag, competitive elution is ineffective. Therefore, we employed native elution via on-resin proteolytic cleavage (e.g. with TEV protease), which specifically releases the target protein into the supernatant. This strategy retains the mVenus tag and potential non-specifically bound proteins attached to the resin, yielding high-purity, non-tagged KEA1 in the supernatant (Fig. 3A and 3C). Following on-column incubation with TEV protease, the eluted KEA1 protein exhibited a distinct shift in mobility to approximately 130 kDa. This observed decrease in molecular weight corresponds to the expected size of the mVenus tag (∼27 kDa), confirming the specific proteolytic cleavage at the TEV site (Fig. 3C). This workflow yielded 3.3 µg of native, tag-free and pure KEA1 protein per g of starting material (ground plant tissue).

These results demonstrate robustness of this cost-efficient workflow, yielding high-purity plant proteins for biochemical characterizations such as structural elucidations, enzymatic activities or immunoprecipitation (IP) experiments.

### Transiently expressed PGDH3 purification via dual-fluorescence tag system

We next evaluated the utility of the novel dual-fluorescence tag vector system (mCherry-TEV-mVenus) for protein production and purification from other plant sources via transient expression in *N. benthamiana* leaves. PGDH3, a key stromal enzyme involved in maintaining chloroplast redox balance through NAD(H)-dependent activity crucial for photosynthesis (Krämer et al., 2024, Höhner et al., 2021), served as a representative test protein.

Transient expression of PGDH3 fused to the mCherry-TEV-mVenus tag was achieved by agroinfiltration of *N. benthamiana* leaves. The protein purification strategy employed was analogous to that developed for stably transformed *A. thaliana* lines (Fig. 3A), utilizing GFP-trap resin to capture the fusion protein via the mVenus moiety (Fig. 4A). A distinct advantage of this dual-tag system is the ability to monitor the elution process in real-time. This is done by observing the fluorescence of the mCherry tag released into the supernatant. Upon incubation with TEV protease, the mVenus portion of the fusion protein remained bound to the GFP-trap resin, while only the PGDH3-mCherry fragment was eluted (Fig. 4A-B). Analysis of the elution kinetics using the TEV protease by SDS-PAGE and tracking the mCherry fluorescence in the supernatant showed that the majority of the PGDH3-mCherry protein was released within the first 2 hours (Fig. 4B-C). Tracking the elution kinetics can considerably shorten incubation times favorable for unstable proteins.

**Figure 4.**
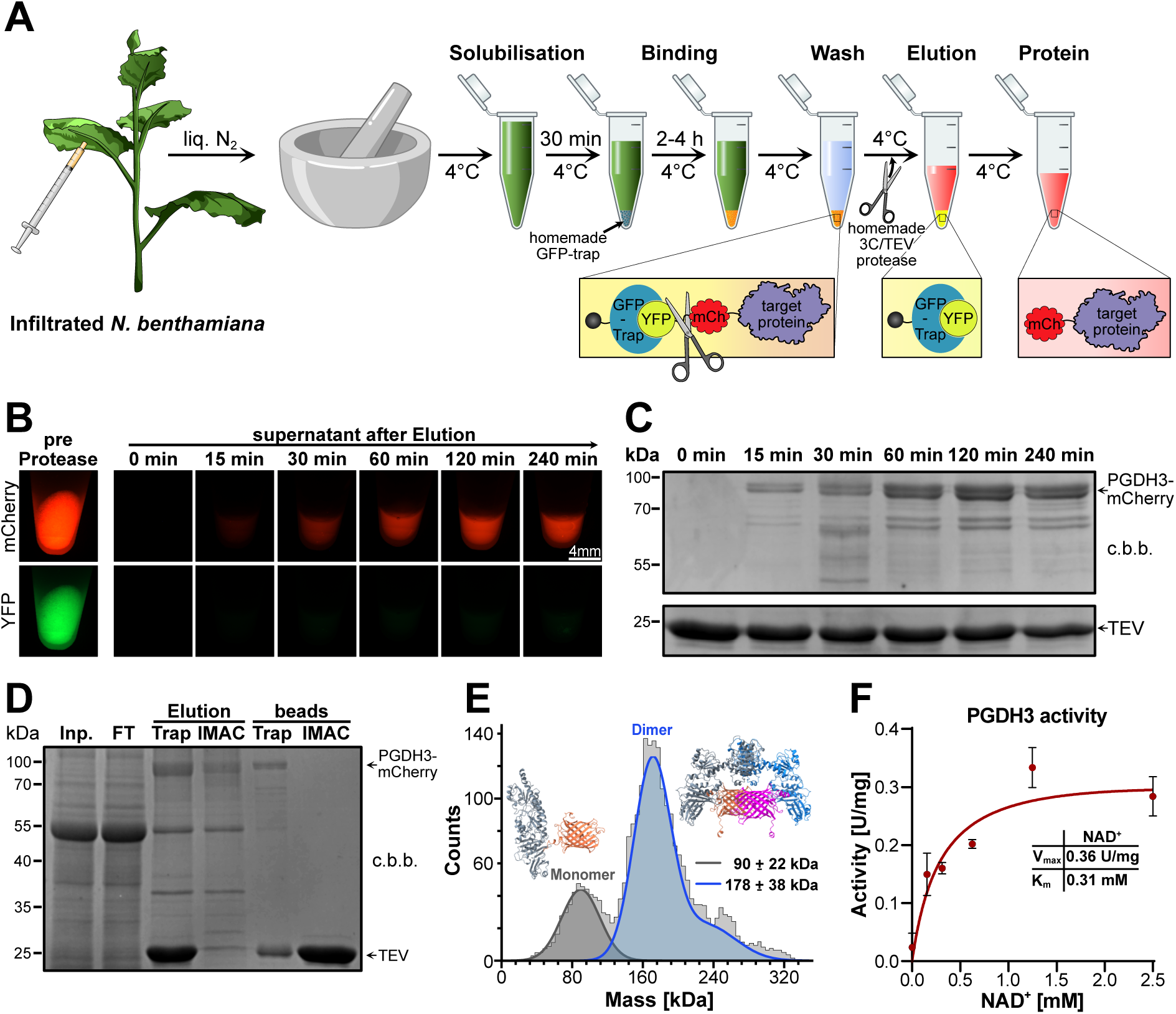
Double-fluorescent tag purification and characterization of PGDH3-mCherry. (**A**) Schematic representation of the “Catch and Release” purification workflow using the double-fluorescent tag. Transiently expressed *N. benthamiana* leaves were ground, solubilized, incubated with GFP-trap resin, followed by multiple wash steps. Protease digestion enabled the continuous release of mCherry-tagged protein. (**B**) Time-resolved elution of PGDH3-mCherry-TEV-mVenus. The left image depicts PGDH3 bound to GFP resin before protease digestion. After enzymatic cleavage, eluted protein (free of resin) was collected at different time points and imaged under a fluorescence stereoscope. (**C**) SDS-PAGE analysis of elution fractions from (B), stained with coomassie brilliant blue (c.b.b.). TEV protease served as a loading control. (**D**) SDS-PAGE analysis illustrating PGDH3-mCherry purification, including removal of His-tagged TEV protease using Ni-NTA affinity chromatography (IMAC). Lanes represent: Total protein lysate (Input, Inp.); unbound fraction after GFP-Trap incubation (Flow-Through, FT); protein eluted from GFP-Trap after TEV cleavage (Elution Trap); final purified protein after IMAC step (Elution IMAC); GFP-Trap resin post-elution (beads Trap); IMAC resin post-elution (beads IMAC). (**E**) Mass distribution histogram of purified PGDH3-mCherry obtained using the isolation protocol. The histogram represents mean trajectory contrasts detected in a dynamic mass photometry analysis (n = 1 movie, 1 min), including trajectories of at least 151 ms in length (n = 2847 trajectories). Predicted 3D structures of both isoforms were generated using AlphaFold 3.0 (Abramson et al., 2024). (**F**) Michaelis Menten kinetics of purified PGDH3-mCherry, assessing phosphoglycerate dehydrogenase function. Data are presented as mean ± SEM (n = 3).

For applications requiring even faster processing, potentially due to protein instability, we show that using the HRV 3C protease together with its recognition motive significantly shortens the required incubation time from ∼120 min (TEV) down to ∼30 min for PGDH3 (Fig. S1E).

A common challenge in protease-assisted protein purification is the co-elution of the protease itself. To address this, we implemented a subsequent purification step using reverse Immobilized Metal Affinity Chromatography (IMAC). Since the TEV protease possesses a His-tag, incubation of the eluted PGDH3-mCherry sample with Ni-NTA resin effectively captured the protease, as shown by SDS-PAGE analysis (Fig. 4D). Alternatively, for proteins with large size differences, size-exclusion chromatography (SEC) or a simple size cut-off spin column can be used to remove the proteases, albeit having other disadvantages (SEC: sample dilution; Spin column: repeated concentrations of proteins increases risk for aggregation). The purification of protease-and tag-free PGDH3 yielded ∼100 µg protein/g ground plant material, i.e. considerably higher amounts compared to the KEA1 isolation.

### One-step purified PGDH3 via Catch & Release is of high quality

As aggregation can be a significant issue during protein purification, we assessed the dispersity and oligomeric state of protease-free PGDH3-mCherry purified via the “Catch & Release” toolbox using mass photometry (Fig. 4E). The results revealed two populations. The major species exhibited an average mass of 178 ± 38 kDa, while a smaller population (approximately one-third of events) averaged at 90 ± 22 kDa. No major protein aggregates at higher molecular weights were detected. Considering the theoretical molecular weight of the PGDH3-mCherry monomer of 87 kDa, these results strongly suggest that the purified protein exists predominantly as a dimer (∼179 kDa), with a smaller fraction present as monomers (∼90 kDa). This observation aligns with previous findings that human PGDH requires dimerization for its enzymatic activity (Xu et al., 2021), suggesting that the transient expression and purification protocol yielded fully functional, non-aggregated, and correctly folded plant PGDH3.

To further confirm the protein quality, we investigated the activity of purified PGDH3-mCherry through an enzymatic assay. The protein exhibited NAD-dependent dehydrogenase activity, consistent with its known function (Benstein et al., 2013). From Michaelis-Menten kinetics, we determined a K_m_ for nicotinamide dinucleotide (NAD^+^) of 0.31 mM (at pH 8.0) and a V_max_ of 0.36 U/mg. Notably, this K_m_ value aligns remarkably well with that reported for Arabidopsis PGDH3 purified from *E. coli* (0.24 mM, pH 8.1) (Benstein et al., 2013), suggesting the protein purified from *N. benthamiana* shares similar kinetic properties. Ultimately, the measured activity provides clear evidence that the PGDH3-mCherry protein, purified using the dual-tag vector following transient expression in *N. benthamiana*, is enzymatically functional.

Taken together, these findings demonstrate that our novel mCherry-TEV-mVenus dual-tag vector enables for the rapid production and purification of substantial amounts of intact, pure and biologically active plant proteins via transient expression, illustrated here with *A. thaliana* PGDH3 expressed in *N. benthamiana*. This approach significantly shortens the experimental timeline, requiring only about one week (∼1-day active work) compared to establishing stable *A. thaliana* lines.

## Conclusions

Here, we developed and validated the “Catch & Release” system, a flexible toolbox accelerating cloning, transgenic plant generation (via FAST), and protein purification in plants. Its modular design features efficient screening markers, diverse and precisely removable tags, including a novel dual-fluorescence option. All modules are easily exchangeable to cater individual needs. We demonstrated *in vivo* functionality of fusion proteins through successful KEA1 complementation in *Arabidopsis* loss-of-function mutants, confirming correct localization and function. Furthermore, by using DIY components we established a robust, low-cost and fast one-step purification workflow yielding tag-free and functional proteins, demonstrated for both membrane (KEA1) and soluble (PGDH3) targets from stable or transient expression systems, respectively.

Purifications times are drastically shortened to hours (from days) while yields (100 ug PGDH3/g plant material) are sufficient for most down-stream applications Overall, Catch & Release provides an efficient and versatile platform for advancing plant protein research. We envision the workflow to be especially useful to rapidly prepare native plant protein isolates for structural analysis by Cryo-EM, IP for interaction studies or enzyme activity measurements.

## Material and Methods

### Plant growth and isolation of *Arabidopsis thaliana* mutant lines

*Arabidopsis thaliana* (ecotype Col-0) plants and *N. benthamiana* plants were grown on soil (Sungro Professional Growing Mix #1, Sun Gro Horticulture, Agawam, MA, USA) under long day conditions (16 h illumination / 8 h dark) at 110 µmol photons m^-2^ s^-1^, 22°C, 50% humidity. Rosettes of 3-week-old Arabidopsis thaliana plants were used for all experiments if not stated differently. Homozygous genotypes were confirmed by PCR using primer combinations depicted in tab. S1.

### Cloning

The *UBQ10* promoter in the pG20_Venus_Hyg vector (Pratt et al., 2020) (Addgene ID: 159703) was first modified by site-directed mutagenesis PCR using primers pGII_UBQ_1 and pGII_UBQ_2 to remove a BbsI restriction site. Subsequently, a synthetic DNA-fragment (Geneart, Thermo) containing a constitutively expressing RFP-cassette flanked by two BbsI Golden Gate sites and a TEV protease site with a downstream NcoI site was inserted via Gibson assembly into the BamHI/XmaI-linearized pG20_updated-UBQ10_Venus_Hyg vector.

To introduce various epitope tags, the pG20_updated-UBQ10_RFP-GoldenGate_TEV_Venus_Hyg vector was linearized with NcoI and SacI (New England Biolabs, Frankfurt, Germany). Flag, Strep, MYC, and HA tags were incorporated by annealing complementary oligonucleotides (listed in tab. S1) and ligating them into the linearized vector. For mCherry insertion, the NcoI site was first removed by amplifying two fragments from the pG20_mCherry_Hyg vector (Pratt et al., 2020) (Addgene ID: 159701) using primers pGII_mCherry_1 to pGII_mCherry_4, followed by Gibson assembly, employing the same strategy used for the epitope tags.

The FAST-Red cassette was amplified from synthetic DNA (Geneart, Thermo) using primers pGII_FAST_1 and pGII_FAST_2, while the pG20_Venus_Hyg vector was amplified with primers pGII_FAST_3 and pGII_FAST_4. Gibson assembly resulted in the pG20_FAST-Red (FR)_Venus vector, which was subsequently digested with HindIII (upstream of the UBQ10 promoter) and EcoRI (downstream of the HSP18.2 terminator) to introduce the updated expression cassette described above. This construct incorporated the UBQ10 promoter, RFP cassette, TEV protease cleavage site, selected tags, and the HSP18.2 terminator. The final plasmids were designated as pG20_TEV_tag_FR.

For the FAST-Green cassette, the pG20_TEV_mVenus_FR vector was amplified using primers pGII_FRed_1 and pGII_FRed_2, while eGFP was amplified using primers pGII_FGreen_1 and pGII_FGreen_2. The two fragments were assembled via Gibson cloning, yielding the pG20_TEV_mVenus_FG vector. Subcloning of all the tag variants followed the same strategy as for the FAST-Red vector, utilizing HindIII and EcoRI restriction sites.

The TEV protease cleavage site in the pG20_TEV_mVenus_FR vector was replaced by digesting the vector with XmaI (New England Biolabs, Frankfurt, Germany) and NcoI, followed by ligation of annealed primers pGII_3C_1 and pGII_3C_1.

### Stable transformation of *Arabidopsis thaliana*

*KEA1* (AT1G01790) genomic DNA (Bölter et al., 2020) and *PGDH3* (AT3G19480) cDNA (Lopez et al., 2022) were amplified without their Stop codon by PCR using primers listed in tab. S1. All constructs described in this paper were generated through classical restriction cloning using BamHI/XmaI restriction sites and the FAST pGreen II system. The *kea1-1kea2-1* mutants were previously described by (Kunz et al., 2014). *A. thaliana* Col-0 plants were transformed using the floral dip method (Clough and Bent, 1998) with the *Agrobacterium tumefaciens* GV3101 strain carrying the additional helper plasmid pSOUP (Hellens et al., 2000). Transgenic seeds were screened for GFP or RFP fluorescence using the Leica M165 FC Fluorescent Stereoscope (Leica, Wetzlar, Germany). Microscopy was carried out in the T_1_ generation. Immunoblotting in fluorescence-selected homozygous F_3_ plants.

### Transient transformation of *N. benthamiana*

For overexpression and immunoprecipitation studies, *N. benthamiana* leaves were infiltrated with *Agrobacterium tumefaciens* strains carrying the respective vectors (listed in tab. S2) as well as co-injected with the 19k vector according to (Waadt et al., 2014). Infiltrated leaves were harvested from plants after 6 days of greenhouse culturing.

### Protoplast isolation and confocal microscopy for protein localization

Protoplast were prepared according to the “Tape-Arabidopsis-Sandwich” method (Wu et al., 2009). Imaging was carried out utilizing a Leica Stellaris 5 Confocal Laser Scanning Microscope (Leica, Wetzlar, Germany), featuring a supercontinuum White Light Laser (WLL) and a 405 nm diode. Emission signals were detected using a Power Hybrid HyDS detector. For protein localization, recombinant mVenus (YFP)-tagged proteins were excited at 514 nm, with emission recorded between 520 and 580 nm. Similarly, mCherry (RFP)-tagged proteins were excited at 585 nm, and emission was detected within the 615–670 nm range. Chlorophyll autofluorescence (chl a) was excited at 405 nm, with emission captured between 623 and 813 nm. Protoplast imaging was performed via Z-stack acquisition, optionally enhanced using the LIGHTNING module, and processed in LAS X software to generate maximum intensity projections.

### Photosynthetic measurements

Photosynthetic performance was evaluated using a Walz Imaging PAM system (Walz GmbH, Effeltrich, Germany) following the methodology outlined by (Kunz et al., 2009). Prior to measurements, plants underwent a 30-min dark acclimation period. A standard induction curve was applied at an irradiance of 110 µmol photons m^-2^ s^-1^ for a duration of 300 sec, with measurements acquired at 20-sec intervals. False-color image analysis and data export were conducted using the Walz ImagingWinGigE software.

### Immunoprecipitation

Three-week-old *A. thaliana* plants or infiltrated *N. benthamiana* leaves were harvested and ground to a fine powder in liquid nitrogen. The tissue was extracted in buffer (50 mM Tris-HCl pH 7.5, 150 mM NaCl, 2.5 mM EDTA, 10% (v/v) glycerol, 0.5% (w/v) Triton X-100/NP-40, 5 mM DTT, protease inhibitors) at a 1:1.5 (w/v) ratio and incubated on a rotation shaker at 4°C for 30 min. Following centrifugation (25,000*g*, 20 min, 4°C), the supernatant was filtered and incubated with homemade GFP-Trap resin for 4 h at 4°C. The resin was washed three times with wash buffer [50 mM Tris-HCl pH 7.5, 150 mM NaCl, 2.5 mM EDTA, 10 % (v/v) glycerol, 0,5 % (w/v) Triton X-100 or NP-40], and bound proteins were cleaved using TEV or 3C protease, with digestion monitored over 4 h. Cleaved proteins were flash-frozen in liquid nitrogen and stored at -80°C for further applications. For immunoblotting and SDS-PAGE, samples were mixed with SDS loading buffer and heated at 80°C for 10 min. Resin-bound fractions were processed similarly. Samples were either stored at -20°C or used immediately.

### Mass photometry

Mass photometry measurements of PGDH3-mCh were performed using a TwoMP instrument (Refeyn Ltd, Oxford, UK). Protein stocks were diluted in PBS (137 mM NaCl, 2.7 KCl, 8.1 mM Na_2_HPO_4_, 1.5 mM KH_2_PO_4_), prior to analysis. Sample well cassettes (Ref: RD501078; Refeyn Ltd, Oxford, UK) were mounted on clean microscope coverslips (Ref: MP-CON-41001; Refeyn Ltd, Oxford, UK). For each measurement, a clean well was pre-filled with 15 µL of PBS for autofocus stabilization. Subsequently, 5 µL of diluted protein solution was added to achieve a final protein concentration of 5 nM. Movies (60 s duration) were recorded using standard instrument settings. Data were acquired using AcquireMP software and analyzed with DiscoverMP software (Refeyn Ltd, Oxford, UK). Each sample was measured at least three times (n ≥ 3). Instrument calibration was performed using a MassFerence P1 Calibrant (Refeyn Ltd, Oxford, UK). Protein structures were simulated using Alphafold 3.0 (Abramson et al., 2024).

### Enzymatic activity assay

PGDH3 activity was assessed in a reaction buffer containing a final concentration of 100 mM Tris-HCl (pH 8.0), 5 mM hydrazine sulfate, 5 mM MgCl_2_ and 0 - 2.5 mM NAD^+^(Höhner et al., 2021). Prior to the reaction, the protein sample was incubated separately at room temperature for 10 min with 20 mM DTT. Each of the three replicates consisted of 200 µl total volume with a final protein concentration of 2.5 ng/µl. The reaction was initiated by adding 3-phosphoglyceric acid (3-PGA) to a final concentration of 10 mM. Absorbance change was recorded at 340 nm using a Spark Multimode Microplate Reader (Tecan, Männedorf, Switzerland). Using the GraphPad Prism Software (v. 10.4.2) absorbance values were fitted according to Michael Menten.

### Immunoblotting

Immunoblotting was performed on total leaf tissue from homozygous *A. thaliana* lines or infiltrated *N. benthamiana* leaves (Völkner et al., 2021, Völkner et al., 2024). Equal amounts of leaf tissue were ground and extracted in solubilization buffer (50 mM Tris-HCl pH 7.5, 150 mM NaCl, 2 mM EDTA, 5 mM DTT, 10% (v/v) glycerol, 0.5% (w/v) Triton X-100, plant protease inhibitor mix) for 30 min at 4°C. Samples were either incubated with the 3C or TEV protease for 1h before mixing with SDS-loading dye and boiling at 95°C for 5 min or directly mixed with SDS-loading dye and boiled. Proteins were separated on 8-15% (w/v) SDS-PAGE gels based on molecular weight and transferred to a PVDF membrane using a wet-blotting system. Immunodetection was performed overnight at 4°C using primary antibodies (listed in tab. S3) followed by chemiluminescent detection with HRP-conjugated secondary antibodies.

### Protease purification

Plasmids encoding TEV or HRV-3C protease (listed in tab. S2) were transformed into *Escherichia coli* BL21 (DE3) and cultured in Terrific Broth (TB) medium supplemented with 50 µg/ml kanamycin at 37°C until the optical density at 600 nm (OD_600_) reached 0.6. Protein expression was induced with 0.5 mM isopropyl β-D-1-thiogalactopyranoside (IPTG), followed by incubation at 18°C for 16 h. Cells were harvested by centrifugation (3,500*g*, 15 min at 25°C), resuspended in buffer A (50 mM Tris-HCl, pH 8.0, 300 mM NaCl, 5% (v/v) glycerol, 2 mM DTT, 20 mM imidazole), flash-frozen in liquid nitrogen, thawed on ice, and lysed using a tissue lyser.

Cell debris were removed by centrifugation (45,000*g*, 30 min at 4°C), and Poly-Histidine-tagged proteases were purified by immobilized metal affinity chromatography (IMAC) using a HisTrap HP (Cytiva Life science, Wilmington, DE, USA) column on an ÄKTA PURE FPLC (Cytiva Life science, Wilmington, DE, USA) system. After washing with buffer A, proteins were eluted using buffer A supplemented with 300 mM imidazole. The buffer was subsequently exchanged to buffer B (50 mM Tris-HCl, pH 8.0, 150 mM NaCl, 5% (v/v) glycerol, 1 mM DTT) using a HiPrep 26/10 Desalting column (Cytiva Life science, Wilmington, DE, USA).

Further purification was performed via size exclusion chromatography (SEC) using a HiLoad 16/16 Superdex 75 pg or 200 pg column (Cytiva Life science, Wilmington, DE, USA) equilibrated in SEC buffer (50 mM Tris-HCl, pH 8.0, 150 mM NaCl, 10% (v/v) glycerol, 1 mM DTT). Fractions were analyzed by SDS-PAGE, pooled, concentrated to 3 mg/ml, aliquoted, flash-frozen in liquid N_2_, and stored at -80°C.

### Expression and purification of the GFP-clamp (GFPc)

The homemade GFP-Trap is based on a double DARPin (GFP-clamp or GFPc) engineered for high-affinity GFP binding (Hansen et al., 2017). The construct was obtained from the Plückthun Laboratory (University of Zurich). The GFPc was subcloned into a pET-28a vector fusing it to an N-terminal and TEV-cleavable His10-Thioredoxin (HSTrx) tag followed by a triple Lysine (KKK) sequence (full-length: 47.8 kDa; cleaved: 32.2 kDa).

*E. coli* Rosetta 2 (DE3) competent cells were transformed with the subcloned GFPc plasmid and cultured in Terrific Broth (TB) medium supplemented with 50 µg/ml kanamycin and 34 µg/ml chloramphenicol at 37°C until an OD_600_ of 0.6 was reached. Protein expression was induced with 1 mM isopropyl β-D-1-thiogalactopyranoside (IPTG), followed by incubation at 20°C for 16 h. Cells were harvested by centrifugation (3,000*g*, 10 min at 25°C), resuspended in buffer A (50 mM Tris-HCl, pH 8.0, 300 mM NaCl, 5% (v/v) glycerol, 2 mM ß-mercaptoethanol, 20 mM imidazole), flash-frozen in liquid nitrogen, thawed on ice, and lysed using a tissue lyser. The lysate was clarified by centrifugation at 30,000*g* for 30 min at 4°C. The soluble supernatant was collected and filtered through a 0.45 µm syringe filter.

Protein purification was performed at 4°C using an ÄKTA FPLC system. Filtered supernatant was applied to a HisTrap HP HP (Cytiva Life science, Wilmington, DE, USA) column equilibrated in buffer A. The column was washed sequentially with 5-10 column volumes (CV) of buffer A, 5 CV of buffer C (50 mM Tris-HCl pH 8.0, 1 M NaCl, 5% (v/v) glycerol, 20 mM imidazole), and finally 5 CV of buffer A, before elution with buffer B (50 mM Tris-HCl pH 8.0, 300 mM NaCl, 5% (v/v) glycerol, 300 mM imidazole). Pooled fractions were buffer-exchanged into PBS (137 mM NaCl, 2.7 KCl, 8.1 mM Na_2_HPO_4_, 1.5 mM KH_2_PO_4_, pH 7.4) using a HiPrep 26/10 Desalting column. The HSTrx tag was cleaved overnight at 4°C using homemade Histidine-tagged TEV protease (1:100 w/w ratio). The cleaved tag and TEV protease were removed by passing the reaction over a second HisTrap HP column equilibrated in PBS; the KKK-GFPc was collected in the flow-through. Final purification was achieved by size exclusion chromatography (SEC) on a HiLoad 16/600 Superdex 200 pg column equilibrated in PBS. Monomeric KKK-GFPc fractions (eluting at ∼85 ml) were pooled and concentrated. Protein concentration was determined by A280 (ε_molar_ = 29.000 M^-1^ cm^-1^). Purified protein was flash-frozen and stored at -80°C.

### Preparation of GFP-Trap Affinity Resin

Purified, tag-cleaved KKK-GFPc was covalently immobilized onto NHS-activated Sepharose 4 Fast Flow resin (Cytiva Life science, Wilmington, DE, USA, Cat# 17090601) via amine coupling, targeting the N-terminal KKK sequence. All steps were performed at room temperature (RT). The vendor-supplied resin slurry was transferred to a conical tube, pelleted by centrifugation (1500*g*, 5-10 min), and the storage solution was discarded. The resin was washed 2 times in 50 ml 1 mM HCl (centrifugation at 1500*g*, 5-10 min) then activated by a 10-minute incubation in 1mM HCl on a rotation shaker. Subsequently the resin was washed twice with Phosphate-Buffered Saline (PBS; 137 mM NaCl, 2.7 mM KCl, 8.1 mM Na_2_HPO_4_, 1.5 mM KH_2_PO_4_, pH 7.4).

Following the final PBS wash and pelleting, the settled resin bed volume was estimated. Purified KKK-GFPc diluted in PBS (1 mg GFPc/ml resin bed volume) was added to the activated resin in a final volume of 50 ml. The mixture was incubated for 2 hours at RT with rotation to facilitate coupling.

After coupling, the resin was pelleted, and the supernatant was removed. To block any remaining active NHS esters, the resin was resuspended in 100 mM Tris-HCl pH 8.5 containing 150 mM NaCl in a total volume of 50 ml and incubated for 1 hour at RT with rotation. The coupled and blocked resin was subsequently washed by sequential resuspension (to approx. 50 ml) and pelleting: twice with PBS containing 1 M NaCl, and once with PBS containing 20% (v/v) ethanol. Finally, the pelleted resin was resuspended in a volume of PBS containing 20% (v/v) ethanol equal to the settled resin bed volume, creating a 50% (v/v) slurry for storage at 4°C. The resulting GFP-trap is stable for at least 2 years.

### Key Materials

| Resource | Source |
| --- | --- |
| Catch & Release vector set | t.b.a. |
| TEV protease | Addgene ID: 171782 |
| 3C protease | Addgene ID: 162795 |
| GFP clamp | (Hansen et al., 2017) |
| Resin for GFP-trap | Cytiva Life Science, Prod. #: 17090601 |

## Accession numbers

The following genetic loci were important to this project: KEA1 (AT1G01790), KEA2 (AT4G00630), PGDH3 (At3g19480). Arabidopsis thaliana (A. thaliana) wild-type and T-DNA insertion mutant plants employed in this study were all from the Columbia 0 (Col-0) accession. The homozygous T-DNA insertion mutant *kea1-1kea2-1* (SAIL_586_ D02, SALK_045324) was published previously (Kunz et al., 2014) and can be obtained from the stock centers under the following IDs: ABRC: CS72318 or NASC: N72318.

## Acknowledgments

We are grateful to Prof. Andreas Plückthun (University of Zurich) for kindly providing GFP-trap components. Thanks to Profs. Mark Howarth (University of Cambridge) and Gottfried Otting (Australian National University) for supplying clones to express and purify TEV and 3C proteases, respectively. We are grateful to Srotoswini Joardar for pilot experiments, Beata Szulc for plant transformation and Benjamin Kouri (all at LMU during their contribution) for help with HRV 3C expression and purification. We thank LMU gardener Albert Schorer for his tremendous support with plant cultivation.

## Author contributions

B.B. and H-H.K. designed the project. S.S. performed most experiments, analyzed data, and prepared figures. S.S. and B.B. designed figures. B.B, S.S and C.E. cloned constructs. C.E. and S.S established transgenic plants. S.M. carried out CLSM, designed the Catch & Release logo, and created the figure icons for tobacco and Arabidopsis. E.W. performed PGDH assays. T.W. helped with mass photometry and carried out KEA1 IP. S.S., B.B., and H-H.K. wrote the manuscript. All authors assisted in editing the manuscript.

---

**Figure S1. Purification and activity assessment of recombinant TEV and HRV 3C proteases.** (**A**) Size exclusion chromatography (SEC) chromatographs of recombinantly expressed tobacco etch virus (TEV) protease purified using a HiLoad 16/60 Superdex 75 pg column. Dashed lines indicate collected fractions, with fraction numbers shown below. (**B**) SDS-PAGE analysis of TEV protease gel filtration fractions stained with Coomassie blue. Fractions 19–23 were pooled, concentrated, and used in subsequent experiments. (**C**) SEC elution profile of recombinantly expressed human rhinovirus (HRV) 3C protease purified using a HiLoad 16/60 Superdex 200 pg column. Dashed lines indicate collected fractions, with fraction numbers shown below. **(D)** SDS-PAGE analysis of HRV 3C protease gel filtration fractions stained with Coomassie blue. Fractions 12–13 were pooled for further analyses. (**E**) Coomassie stained SDS-PAGE of comparison of cleavage efficiency between homemade TEV and HRV 3C proteases over a 120-minute incubation with PGDH3-mVenus bound to homemade GFP-trap resin. Both proteases were used at a final concentration of 12 µM.

**Figure S2. Step-by-step preparation of the homemade GFP-trap resin** Sepharose 4 Fast Flow resin was activated (2x 1 mM HCl; 1x PBS) and conjugated with purified KKK-GFP-clamp via N-terminal amine coupling to a target density of 1 mg/mL settled resin. Residual active sites were blocked using Tris-HCl buffer, followed by sequential high-salt (1 M NaCl) and ethanol (20% v/v) washes. The final product was stored at 4°C as a 50% (v/v) slurry in PBS containing 20% (v/v) ethanol.

