## Supplementary Figures for "Catch & Release – rapid cost-effective protein purification from plants using a DIY GFP-Trap-protease approach"

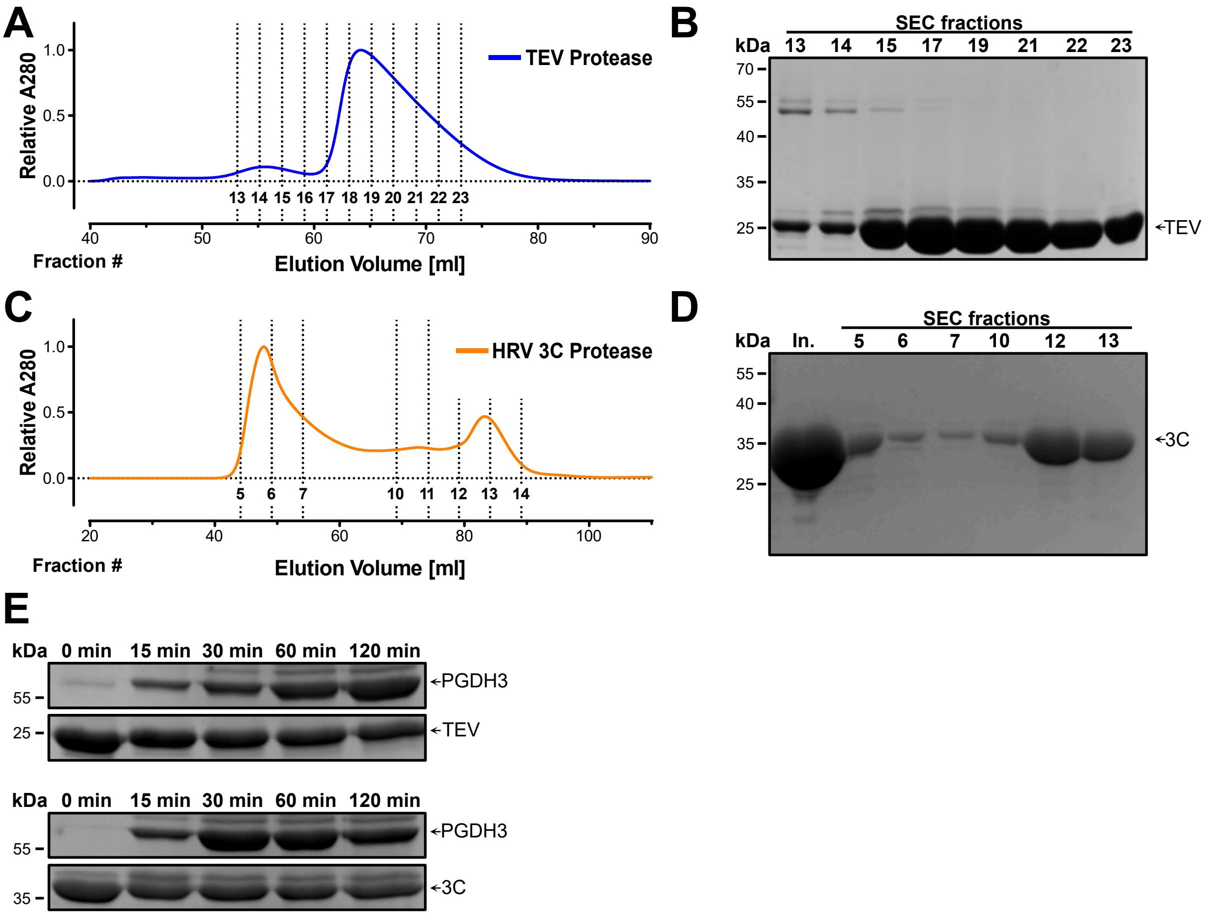


**Figure S.1. Purification and activity assessment of recombinant TEV and HRV 3C proteases.**

(**A**) Size exclusion chromatography (SEC) chromatographs of recombinantly expressed tobacco etch virus (TEV) protease purified using a HiLoad 16/60 Superdex 75 pg column. Dashed lines indicate collected fractions, with fraction numbers shown below. (**B**) SDS-PAGE analysis of TEV protease gel filtration fractions stained with Coomassie blue. Fractions 19–23 were pooled, concentrated, and used in subsequent experiments. (**C**) SEC elution profile of recombinantly expressed human rhinovirus (HRV) 3C protease purified using a HiLoad 16/60 Superdex 200 pg column. Dashed lines indicate collected fractions, with fraction numbers shown below. **(D)** SDS-PAGE analysis of HRV 3C protease gel filtration fractions stained with Coomassie blue. Fractions 12–13 were pooled for further analyses. (**E**) Coomassie stained SDS-PAGE of comparison of cleavage efficiency between homemade TEV and HRV 3C proteases over a 120-minute incubation with PGDH3-mVenus bound to homemade GFP-trap resin. Both proteases were used at a final concentration of 12 μM.

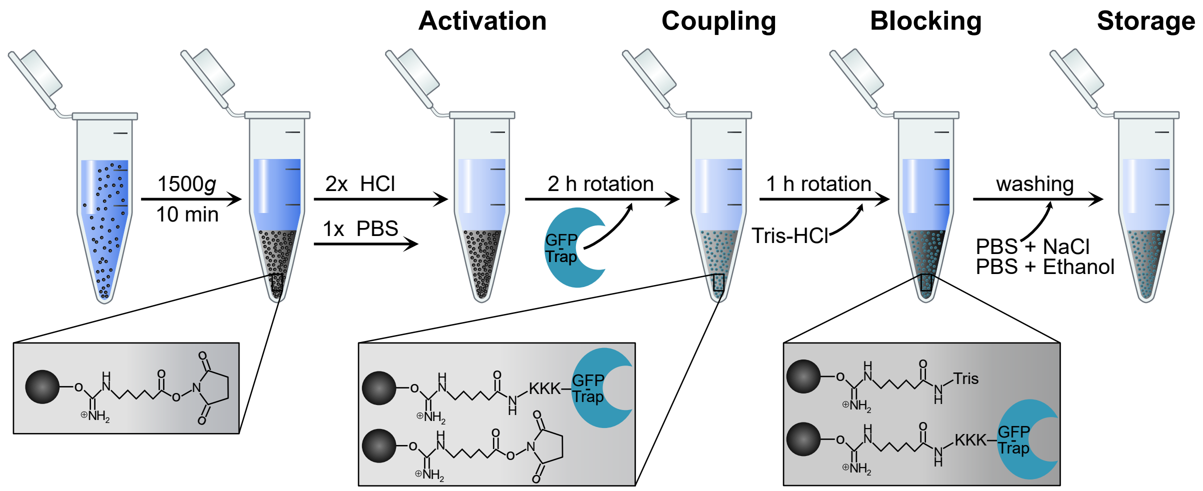


**Figure S2. Step-by-step preparation of the homemade GFP-trap resin**

Sepharose 4 Fast Flow resin was activated (2x 1 mM HCl; 1x PBS) and conjugated with purified KKK-GFP-clamp via N-terminal amine coupling to a target density of 1 mg/mL settled resin. Residual active sites were blocked using Tris-HCl buffer, followed by sequential high-salt (1 M NaCl) and ethanol (20% v/v) washes. The final product was stored at 4°C as a 50% (v/v) slurry in PBS containing 20% (v/v) ethanol.
