## Supplementary Tables for "Catch & Release – rapid cost-effective protein purification from plants using a DIY GFP-Trap-protease approach"

Table S1: Primers used in this study

| primer name | sequence |
|---|---|
| **Genotyping** | |
| WT_kea1_fwd | ccctcaaactcctacaatttctatg |
| WT_kea1_rev | gcaattattgcagtaatagccactgc |
| tDNA_kea1_rev | tagcatctgaatttcataaccaatctcgatacac |
| WT_kea2_fwd | gttgctatcactggcataattgc |
| WT_kea2_rev | ggatcaatggacatgccccac |
| tDNA_kea2_rev | attttgccgatttcggaac |
| **KEA1 and PGDH3 constructs** | |
| Kea1_gDNA_fwd | tttagtggatccatggagtatgcgtctacttttcaaagg |
| Kea1_gDNA_rev | tttcccggggattacgactgtgcctccttcga |
| Pgdh3_cDNA_fwd | tttagtggatccatggcgacgtctctgaatctatc |
| Pgdh3_cDNA_rev | tttcccgggtagtttgaggaaaacaaactcttcaatgg |
| **Catch & Release vector** | |
| pGII_UBQ_1 | gaagtcgtggttggaacgacttctttttccacg |
| pGII_UBQ_2 | gaagtcgttccaaccacgacttcaaagcaag |
| pGII_Flag_1 | catgggcggaggatctgactacaaggacgacgatgacaagtaggagct |
| pGII_Flag_2 | cctacttgtcatcgtcgtccttgtagtcagatcctccgcc |
| pGII_Strep_1 | catgggtgggtccagcgcatggtcacatccgcagtttgaaaaaggtgg |
| pGII_Strep_2 | accaccacctttttcaaactgcggatgtgaccatgcgctggacccacc |
| pGII_Strep_3 | tggtagcggtggtggttcaggtggtagtgcttggagccatcctcagttcgagaaataggagct |
| pGII_Strep_4 | cctatttctcgaactgaggatggctccaagcactaccacctgaaccaccaccgct |
| pGII_MYC_1 | catgggcggaggatcttctgaacaaaagttgatttcagaagaagatctgtaggagct |
| pGII_MYC_2 | cctacagatcttcttctgaaatcaacttttgttcagaagatcctccgcc |
| pGII_HA_1 | catgggcggaggatcttacccatacgatgttccagattacgcttaggagct |
| pGII_HA_2 | cctaagcgtaatctggaacatcgtatgggtaagatcctccgcc |
| pGII_mCherry_1 | ttccaaggctccatggtgagcaagggc |
| pGII_mCherry_2 | cagcccatcgttttcttctgcattac |
| pGII_mCherry_3 | gtaatgcagaagaaaacgatgggctggg |
| pGII_mCherry_4 | catcttcatcttcatatgagctcctacttgtacagc |
| pGII_FAST_1 | acgttatcactaaatggagcaacctac |
| pGII_FAST_2 | aagctttataatgtcgcggaac |
| pGII_FAST_3 | gcgacattataaagcttcgacgagtc |
| pGII_FAST_4 | gatatattgtggtgtaacgttatcactaaatg |
| pGII_FRed_1 | agtagtgtgctggccaccacg |
| pGII_FRed_2 | tgagcttaccccactgatgtcat |
| pGII_FGreen_1 | atcagtggggtaagctcacttgtacagctcgtc |
| pGII_FGreen_2 | ggccagcacactactatggtgagcaaggg |
| pGII_3C_1 | ccgggagtctggaagttctgttccaggggccctc |
| pGII_3C_2 | catggagggcccctggaacagaacttccagactc |

Table S2: Vectors used in this study

| Plasmid Name | Source | Identifier |
|---|---|---|
| pG20_KEA1_TEV_mVenus_FAST-Red | This paper | N/A |
| pG20_KEA1_3C_mVenus_FAST-Red | This paper | N/A |
| pG20_KEA1_TEV_mVenus_FAST-Green | This paper | N/A |
| pG20_KEA1_TEV_mCherry_FAST-Red | This paper | N/A |
| pG20_KEA1_mCherry_TEV_mVenus | This paper | N/A |
| pG20_KEA1_TEV_Strep_FAST-Red | This paper | N/A |
| pG20_KEA1_TEV_Flag_FAST-Red | This paper | N/A |
| pG20_KEA1_TEV_MYC_FAST-Red | This paper | N/A |
| pG20_KEA1_TEV_HA_FAST-Red | This paper | N/A |
| pG20_PGDH3_TEV_mVenus_FAST-Red | This paper | N/A |
| pG20_PGDH3_3C_mVenus_FAST-Red | This paper | N/A |
| pG20_PGDH3_mCherry_TEV_mVenus_FAST-Red | This paper | N/A |
| pG20_Venus_Hyg | Addgene | 159703 |
| pG20_mCherry_Hyg | Addgene | 159701 |
| pET28-MBP-super TEV protease | Addgene | 171782 |
| pet-NT*-HRV3CP | Addgene | 162795 |

Table S3: Reagents and resources used in this study

| REAGENT or RESOURCE | SOURCE | IDENTIFIER |
|---|---|---|
| **Antibodies** | | |
| α-GFP | Roche | Prod. #: 11814460001 |
| α-mCherry | Abcam | Prod. #: AB213511 |
| α-Strep | Agrisera | Prod. #: AS21 4682 |
| α-Flag | Sigma-Aldrich | Prod. #: A8592-.2MG |
| α-MYC | Agrisera | Prod. #: AS21 4685 |
| α-HA | Agrisera | Prod. #: AS18 4176 |
| **Bacterial and virus strains** | | |
| Agrobacterium tumefaciens GV3101 competent cells (Hellens et al., 2000) | | N/A |
| *Escherichia coli* Bl21 (DE3) competent cells | ThermoFisher | Prod. #: EC0114 |
| *Escherichia coli* DH5α competent cells | ThermoFisher | Prod. #: EC0112 |
| **Chemicals, peptides, and recombinant proteins** | | |
| Cellulase “Onozuka R-10” | Serva | Cat. No. / ID: 16419 |
| Macerozyme R-10 | Serva | Cat. No. / ID: 28302 |
| **Experimental models: Organisms/strains** | | |
| Arabidopsis thaliana, Columbia-0, CS70000 | European Arabidopsis Stock Centre (NASC) | NASC ID: N70000 |
| Arabidopsis (Col-0) *kea1-1kea2-1* (Kunz et al., 2014) | | N/A |

| | | |
|---|---|---|
| Arabidopsis (Col-0) *kea1-1kea2-1* pUBQ10:KEA1-TEV-mVenus_FAST-Red | This paper | N/A |
| Arabidopsis (Col-0) *kea1-1kea2-1* pUBQ10:KEA1-3C-mVenus_FAST-Red | This paper | N/A |
| Arabidopsis (Col-0) *kea1-1kea2-1* pUBQ10:KEA1-TEV-mVenus_FAST-Green | This paper | N/A |
| Arabidopsis (Col-0) *kea1-1kea2-1* pUBQ10:KEA1-TEV-mCherry_FAST-Red | This paper | N/A |
| Arabidopsis (Col-0) *kea1-1kea2-1* pUBQ10:KEA1-mCherry-TEV-mVenus_FAST-Red | This paper | N/A |
| **Oligonucleotides** | | |
| all oligonucleotides used in this study are listed in table S1 | N/A | N/A |
| **Recombinant DNA** | | |
| all vectors used in this study are listed in table S2 | N/A | N/A |
| **Software and algorithms** | | |
| GraphPad Prism (v10.3.0) | N/A | https://www.graphpad.com/features |
| ChimeraX (v1.7.1) (Meng et al., 2023) | | https://www.cgl.ucsf.edu/chimerax/ |
| Fiji (Schindelin et al., 2012) | | https://imagej.net/software/fiji/ |


HELLENS, R. P., EDWARDS, E. A., LEYLAND, N. R., BEAN, S. & MULLINEAUX, P. M. 2000. pGreen: a versatile and flexible binary Ti vector for Agrobacterium-mediated plant transformation. *Plant Mol Biol,* 42**,** 819-32.

KUNZ, H.-H., GIERTH, M., HERDEAN, A., SATOH-CRUZ, M., KRAMER, D. M., SPETEA, C. & SCHROEDER, J. I. 2014. Plastidial transporters KEA1, -2, and -3 are essential for chloroplast osmoregulation, integrity, and pH regulation in Arabidopsis. *Proceedings of the National Academy of Sciences,* 111**,** 7480-7485.

MENG, E. C., GODDARD, T. D., PETTERSEN, E. F., COUCH, G. S., PEARSON, Z. J., MORRIS, J. H. & FERRIN, T. E. 2023. UCSF ChimeraX: Tools for structure building and analysis. *Protein Science,* 32**,** e4792.

SCHINDELIN, J., ARGANDA-CARRERAS, I., FRISE, E., KAYNIG, V., LONGAIR, M., PIETZSCH, T., PREIBISCH, S., RUEDEN, C., SAALFELD, S., SCHMID, B., TINEVEZ, J.-Y., WHITE, D. J., HARTENSTEIN, V., ELICEIRI, K., TOMANCAK, P. & CARDONA, A. 2012. Fiji: an open-source platform for biological-image analysis. *Nature Methods,* 9**,** 676-682.
